# Estimates of abundance and change in abundance of the Indo-Pacific bottlenose dolphins (*Tursiops aduncus*) along the south coast of South Africa

**DOI:** 10.1101/2019.12.13.875179

**Authors:** OA Vargas-Fonseca, SP Kirkman, WC Oosthuizen, T Bouveroux, V Cockcroft, D Conry, PA Pistorius

**Author notes:** Corresponding author, (OAVF). Dauphin Island Sea Lab, University of South Alabama, Alabama, United States (T Bouveroux).

## Abstract

We investigated the abundance of Indo-Pacific bottlenose dolphins (*Tursiops aduncus*) along the south coast of South Africa, from the Goukamma Marine Protected Area (MPA) to the Tsitsikamma MPA, between 2014 and 2016. During this period, 662.3h of boat-based photo-identification survey effort was carried out, and the sighting histories of 817 identified individuals were used to estimate abundance using mark-recapture modelling. The selected open population model (POPAN) provided an estimate of 2,295 individuals (95% CI: 1,157-4,553) for the entire study area. A model estimate was produced for a subset of the study area, Plettenberg Bay, which could be compared with a past estimate for this location (2002-2003). The comparison suggested a 72.3% decrease in abundance, from 6,997 (95% CI: 5,230-9,492) in 2002-2003 to 1,940 (95% CI: 1,448-2,600) in 2014-2016. The decline in abundance was supported by a 72% reduction in mean group size for Plettenberg Bay between the periods. It is essential to be able to assess abundance changes at other locations to inform revision of *T. aduncus* conservation status in South Africa.

## Introduction

Information on the abundance and trends of wildlife populations is essential for species and ecosystem conservation management strategies [1,2].Trends in abundance provide feedback on the success or failure of implemented conservation strategies and indicate natural or anthropogenic driven ecosystem changes [3]. In both terrestrial and marine ecosystems, predator population trends are thought to integrate the state of lower trophic levels and the physical environment that they inhabit [4,5]. For this reason predator population trends are often considered to be good indicators of ecosystem health.

The escalating human population, with disproportionately higher growth rates in coastal areas, is exerting increased pressure on coastal ecosystems and marine species. Coastally distributed dolphin species are highly susceptible to current and future human-related threats such as habitat degradation from pollution and costal development (e.g., harbours and offshore wind farms), competition with fisheries, and bycatch in fishing gear or shark exclusion nets [6]. Examples of inshore dolphin species that of current conservation concern and which face a multitude of threats include the vaquita (*Phocoena sinus*), humpback dolphins (*Sousa spp)* [7–9], Australian snubfin dolphins (*Orcaella heinsohni)* [10] and Hector’s dolphins (*Cephalorhynchus hectori*) [11]. For such species, studies that document population size and trends are essential for conservation and management planning [12].

The Indo-Pacific bottlenose dolphin (*Tursiops aduncus)* has been listed as a Data Deficient species by the IUCN Red List of Threatened Species since 1996 [13]. Their distribution is apparently continuous along coastal areas (including mid-ocean island shores) in the Indian Ocean, from False Bay (South Africa) eastwards right through to the Solomon Islands and New Caledonia in the western Pacific Ocean [14] including the east and west coasts of Australia and the south-east Asian waters [15]. The most recent South African Red List conservation assessment [16] recognized three sub-populations of *T. aduncus* in South African waters based on previous genetic studies [17] (Fig 1). A resident sub-population in northern KwaZulu-Natal (between Kosi Bay and Ifafa) was classified as Vulnerable; a migratory sub-population that is thought to move between Plettenberg Bay and Durban as Data Deficient; and a resident sub-population south of Ifafa with its western limit at False Bay as Near Threatened [16]. Research priorities identified by the conservation assessment [16] include (amongst others) conducting research into their population genetics to determine significant management units, assessing the effectiveness of Marine Protected Areas (MPA) in addressing conservation needs of sub-populations, and determining abundance estimates throughout their range as well as site specifically [16]. A subsequent genetic study [18,19] defined two conservation units (instead of three sub-populations) along the South African Coast: one along the Natal Bioregion and another in the Agulhas Bioregion (Fig 1). The results from genetic population structure analysis thus refutes the existence of a migratory sub-population [17] as described in the latest conservation assessment [16].

**Fig 1:**
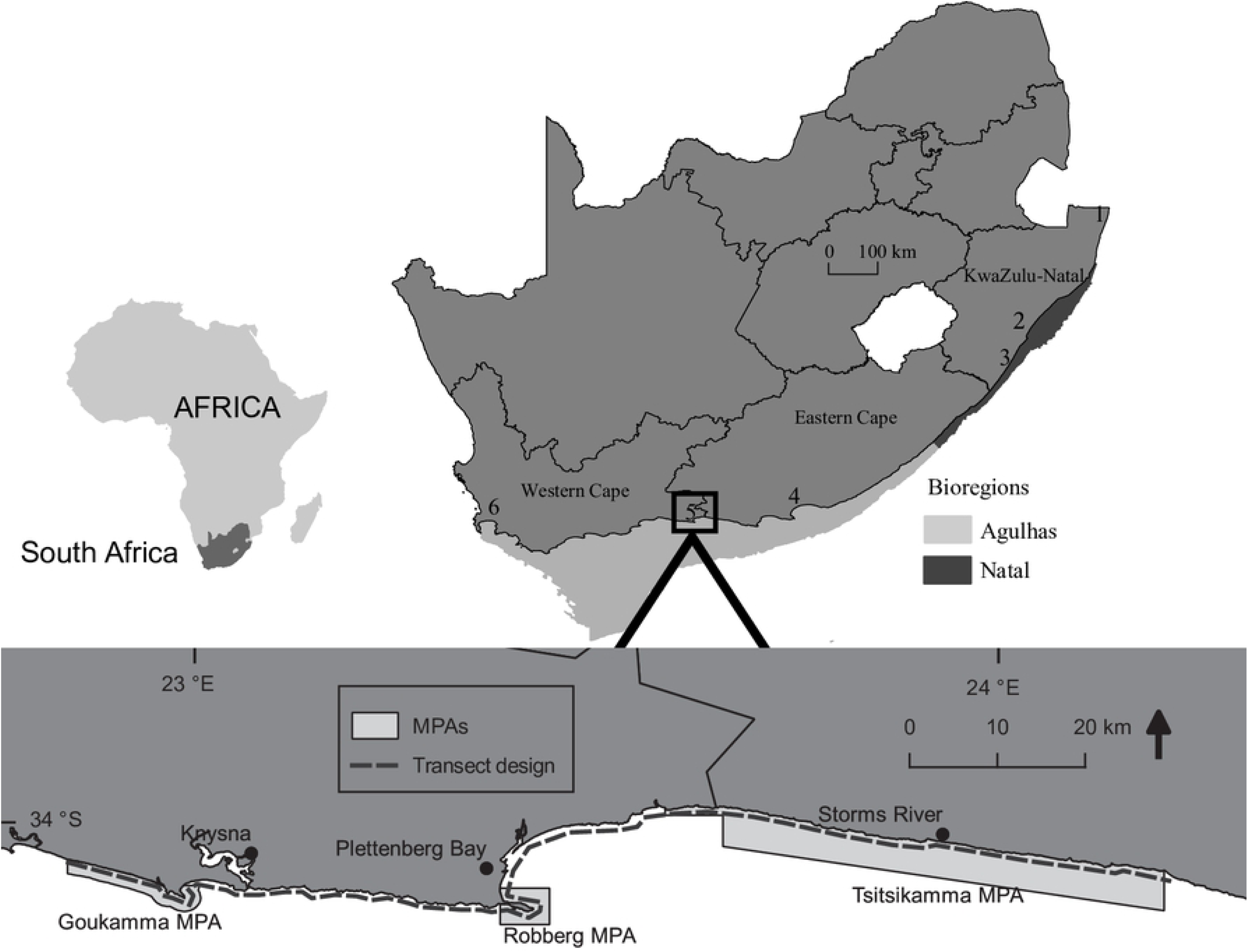
Map of South Africa with relevant locations mentioned in the text. (1) Kosi Bay; (2) Durban; (3) Ifafa; (4) Algoa Bay; (5) study area; (6) False Bay. The study area extended from the western boundary of Goukamma to the eastern boundary of the Tsitsikamma MPA. Boat surveys were conducted parallel to the coast (dashed black line).

The abundance and changes in population numbers of *T. aduncus* along South Africa’s coast is poorly understood; estimates of numbers are restricted to localised areas (summarized in 16) and data on changes in population numbers are non-existent. For the sub-population in the Agulhas Bioregion, only two mark-recapture abundance estimates are available: one in Algoa Bay (1991-1994) where 28,482 (95% CI: 16,220-40,744) individuals were estimated [20] and another for Plettenberg Bay (2002-2003) where 6,997 (95% CI: 5,230-9,492) individuals were estimated [21]. Results from these studies showed that numerous individuals were utilising both areas, indicating a dynamic population on the south coast of South Africa with long-range movements [20].

This study estimates *T. aduncus* population abundance and group sizes along 145 km of coastline in the Agulhas Bioregion off the south coast of South Africa. The data were obtained using boat-based surveys and mark-recapture methods. Furthermore, for a subset of the study area (Plettenberg Bay; 29 km of coastline), separate population abundance and group size estimates were determined so that it could directly be compared with a study conducted in this area more than ten years previously (2002-2003). Tourism is an important revenue along the Bitou municipality which includes Plettenberg Bay [22]. The latter is a growth centre for marine tourism activities including boat-based marine mammal viewing, fishing charters and adventure rides that can potentially disturb dolphins. This is the first attempt at assessing change abundance of a *T. aduncus* population over time at any location in South Africa. We hypothesized that dolphin numbers and group sizes would have decreased since the first assessment due to increasing human activities in the coastal zone.

## Methods

### Study area, survey design and data collection

Data were collected during standardized boat surveys along 145 km of coastline within the Agulhas Bioregion, between the western border of the Goukamma MPA and the eastern boundary of the Tsitsikamma MPA on the south coast of South Africa (Fig 1). Ninety-seven kilometres of the coastline of the study area is within MPAs, namely the Goukamma, Robberg and Tsitsikamma MPAs. There are two main dolphin hotspots in this area, namely the Goukamma MPA and the Plettenberg Bay area [23]; both areas are characterized by sandy shores and gentle slopes. The stretch between Goukamma to Robberg MPA and Tsitsikamma MPA is largely uninhabited (by humans) with exposed rocky coasts and steeper gradients.

The surveys were designed as a transect line running parallel to the coast. Bi-monthly boat surveys were conducted between March 2014 and February 2016. At least two experienced observers were present during surveys which were performed at a constant speed of approximately eight knots (see [23] for further information on study area, survey design and data collection procedures).

In this study, digital dorsal fin photo-ID images were taken using a Nikon SLR camera equipped with a Tamron 300 or 600 mm lens. The dorsal fins of as many dolphins as possible were photographed from both sides (if possible), without any preference towards individuals with obvious markings [1]. Group sizes were estimated independently as minimum, maximum and best estimates, with best estimates not necessarily being the mean of the upper and lower estimates [24]. A group was defined as two or more animals within a 100-m radius of each other, showing similar behaviour [25]. Survey effort was measured as the number of hours travelled in good sighting conditions (Beaufort scale ≤ 3). Survey effort was discontinued when conditions exceeded Beaufort scale 3.

### Data processing and analysis

#### Photo-identification catalogue and data selection

Dorsal fin images were cropped and graded according to the photo quality (Q) and distinctiveness (D). Quality were scored from 1 to 3 (Q1 being excellent quality and Q3 poor quality). The Q grade was based upon photo clarity, contrast, angle, portion of frame filled by the fin, angle, exposure, water spray and the percentage of the fin image that is visible in the frame (adapted from [1,26,27]. Photographs graded Q1 were therefore well exposed, without water droplets, in sharp focus, with the dorsal fin orientated perpendicular to the photographer and occupying a large proportion of the frame (adapted from [1]). Using only photographs graded Q1-Q2, the fins were then graded according to the fin distinctiveness (D). Distinctiveness was graded from 1 to 3 (D1 very distinctive and D3 no distinctive characteristics). Photographs with distinctiveness grades D1-D2 were catalogued according to the location of the most prominent or distinguishing feature. The categories included: leading edge, mutilated, peduncle and trailing edge; with the latter subdivided into entire, low, mid or upper third (adapted from [26]). As many features as possible were used to confirm matches and to reduce the possibility of false positives only long lasting markings were considered [1]. Two different experienced researchers visually compared photographs from each category to avoid misidentification of individuals (first within the same category and subsequently between categories where required).

#### New identifications and discovery curve

To evaluate whether the population had been sampled comprehensively, the cumulative number of newly identified individuals was plotted over time in a discovery curve. If a discovery curve reaches an asymptote, this indicates that the whole population has been identified and that it is likely to be a closed population with no immigration or emigration (e.g. [1]). The discovery curve of an open population (births, deaths, immigration or emigration occurs) is not likely to reach an asymptote (e.g. [20]).

#### Mark-recapture analysis

Open and closed population models were fitted using the software MARK 8.2 [28] to estimate the population size of *T. aduncus* in the study area. Only high quality photographs (Q ≤ 2) were used to construct encounter histories for all the identified individuals (D ≤ 2) using calendar month as capture occasions.

Open population estimates were obtained using the POPAN parameterization [29], which calculates the super-population size 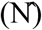, apparent survival probability (Φ), capture probability (p), and the probability of immigration or entry (b) from the super-population to the local population present in the study area. Demographic parameters were designated as time dependent (t), constant over time (.) or seasonal (s), whereas capture probability were additionally allowed to vary with survey effort. Seasons were defined as the austral winter (May-October) or summer (November-April) [30,31]. The most parsimonious model was selected using Akaike’s Information Criterion for small sample sizes (AICc) [32]. Monthly survival probabilities estimated by the model were transformed to annual survival probability with associated variances re-scaled using the Delta method [33].

Goodness-of-fit (GOF) of the fully time-dependent Cormack-Jolly-Seber (CJS) model was assessed in program RELEASE to verify whether the encounter history data met model assumptions [34], A variance inflation factor (ĉ) was calculated based on the results of Test 2 + Test 3 in order to determine if the data were over-dispersed (ĉ > 1) or under-dispersed (ĉ < 1) and to evaluate the need for an adjusted model selection criterion (i.e. quasi-Akaike Information Criterion, QAICc). Test 2 determines capture homogeneity; Test 3 homogeneous survival probability; Test 3 SR presence of transience in the data; and Test 3Sm effect of capture on survival [34].

We used closed population models to compare abundance estimates for data collected during 2014 to 2016, to previous Plettenberg Bay abundance estimates [21]. This was because the past estimates were based only on closed models and the data were not available for re-analysis. In the earlier study closed models were fitted using the program CAPTURE in MARK [35]. The model selection was based upon model selection criteria values produced by the program CAPTURE [36]. The higher the selection criteria the better the model fits (larger value 1.0) and selection values lower than 0.75 should not be used to estimate abundance [35]. Presently, CAPTURE is considered to be an outdated programme for estimating abundance; for this reason the closed population models were also estimated in MARK. Huggins’ model were set as p=c, where the initial capture probability (p) is equal to the recapture probability (c). These settings were used because the animals were not physically captured and a behavioural response to capture was not expected.

#### Estimating super-population size

The mark-recapture abundance estimates refer to the number of marked individuals in the population. To estimate the super-population size of *T. aduncus*, the mark-recapture results were scaled up according to the proportion of marked individuals in good quality photos (≤Q2) [1,26]. The proportion of marked individuals in the population was estimated from the ratio of distinctive individuals (D1 + D2) to the total sample (D1 + D2 + D3) [1,26,37]. The super-population size was estimated as:

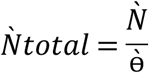

where 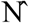 total is the estimated abundance, 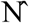 is the mark-recapture estimate of the number of animals with long-lasting marks, and 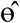 is the estimated proportion of animals with long lasting marks in the population [1]. The variance estimate was calculated using the delta method:

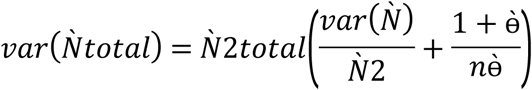

where n is the total number of animals from which θ was estimated [1]. Confidence intervals for 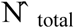 assumed that the error distribution was the same as that of the mark-recapture estimates of the marked population [1].

## Results

In total, 662.3 h of survey effort were conducted over 189 surveys and145 days from March 2014 to February 2016. *T. aduncus* were encountered throughout the year, Average group size was estimated as 47 ± 55 (mean ± SD) individuals, with larger group sizes during winter (57 ± 63) compared to summer (35 ± 42; Table 1). For Plettenberg Bay only, the mean group size was 26 ± 26, which is 78% lower than in 2002-2003 (Table 1).

**Table 1:** *T. aduncus* group size statistics for the entire research area, and for Plettenberg Bay only. Also included for comparison are past estimates for Plettenberg Bay (2002-2003)

|  | Summer | Winter | Overall |
| --- | --- | --- | --- |
| Entire study area 2014-2016 |  |  |  |
| Mean ± SD | $35 \pm 42$ | $57 \pm 63$ | $47 \pm 55$ |
| Range | 1-300 | 1-350 | 1-350 |
| Median | 20 | 40 | 30 |
| Plettenberg Bay 2014-2016 |  |  |  |
| Mean ± SD | $26 \pm 28$ | $26 \pm 18$ | $26 \pm 26$ |
| Range | 1-100 | 3-65 | 1-100 |
| Median | 15 | 23 | 18 |
| Plettenberg Bay 2002-2003 [21] |  |  |  |
| Mean ± SD | $124 \pm 111$ <sup>1</sup> | $82 \pm 143$ <sup>1</sup> | $120 \pm NA$ <sup>3</sup> |
| | $211 \pm 139$ <sup>2</sup> | $56 \pm 76$ <sup>2</sup> | |
| Range | NA | NA | 2-500 <sup>3</sup> |
| Median | NA | NA | 80 <sup>3</sup> |

'NA': not available; <sup>1</sup> in 2002; <sup>2</sup> in 2003; <sup>3</sup> in 2002-2003.

A total of 80.6 h was spent with *T. aduncus* groups during surveys and 10,431 dorsal fin photographs were taken and assessed for quality. Of 4,015 photographs found to be of acceptable quality (≤ Q2), 2,274 photographs had individuals with sufficient distinctiveness (≤ D2). The final catalogue consisted of 817 identified animals with a total of 1,558 photos (which includes multiple good photos per individual per sighting). The proportion of identifiable individuals (adults and juveniles) was 0.77. Of the identified animals, 72.7% were encountered only once, 16.8% were encountered twice, 6.2% were encountered three times and 4.3% were encountered between 4 and 7 times in the entire study area.

The discovery curve never reached an asymptote (Fig 2). New individuals were thus still being identified towards the end of the study period, suggesting either that the population is open or that not all individuals of a closed population had been identified.

**Fig 2:**
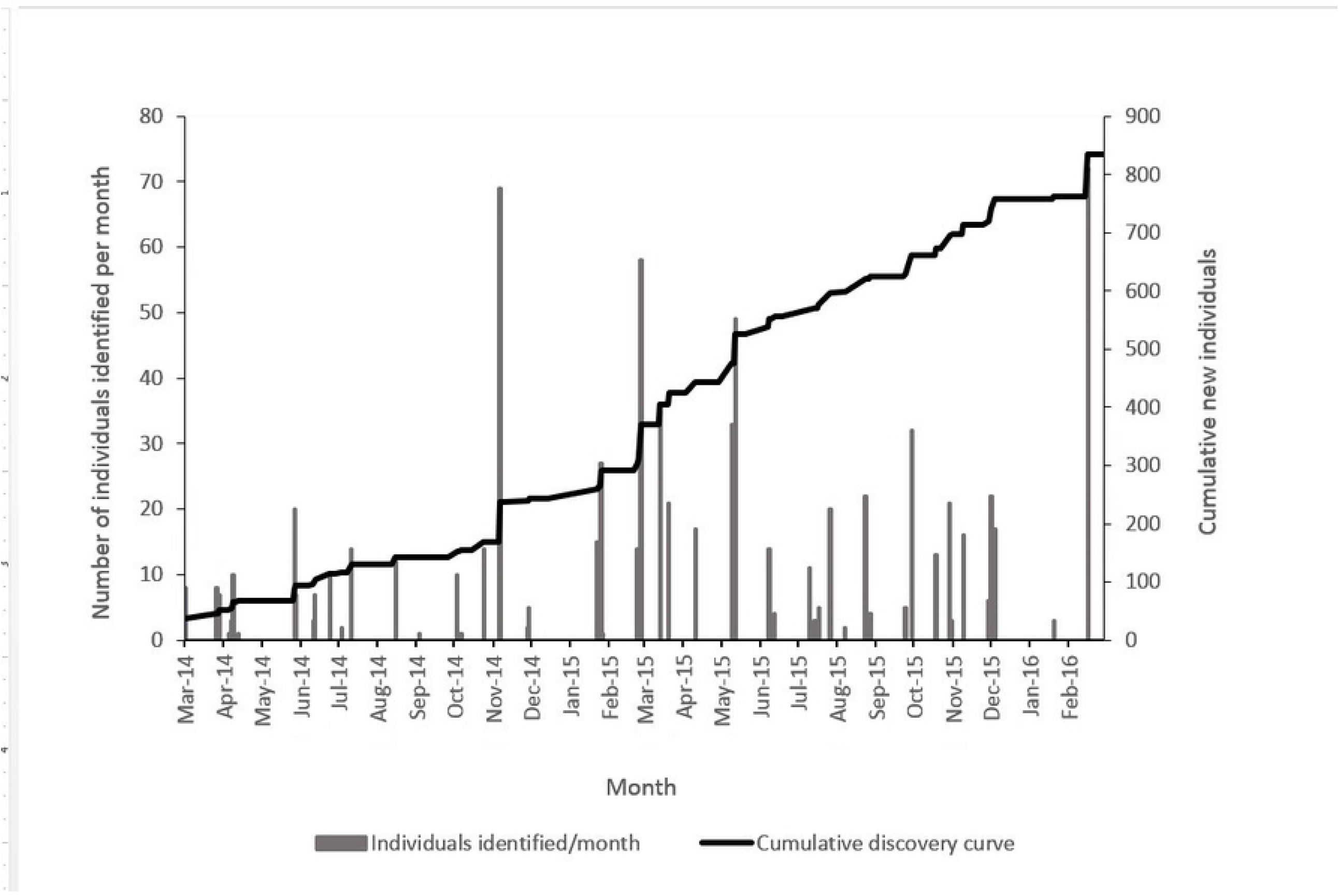
Number of *T. aduncus* identified from photographs, and the cumulative discovery curve for new individuals.

### Abundance estimates

#### Open population model

Goodness-of-fit results (Table in S1 Table) indicated that there was over-dispersion in the encounter history data summarising the observations made in the entire research area, with a variance inflation factor of ĉ = 1.71. Goodness of fit tests suggested there was heterogeneity in capture probabilities between individuals, and that transient animals (permanent emigration after a single encounter) were present. The most parsimonious POPAN model for the entire area assumed constant survival, time dependent capture probability, a seasonal (summer and winter) probability to enter the local population from the super-population, and a constant super-population size (Table in S2 Table). The model produced a super-population size of 2,295 (SE: 827; 95% CI: 1,157-4,553). The annual survival was estimated to be 0.87 (± 0.12).

#### Closed population model

The most appropriate model for Plettenberg Bay (2014-2016) had capture probability as time-dependent. The model M_t_ produced an abundance estimate of 1,063 (SE: 125, 95% CI: 858-1,360) marked individuals which translates to a super-population size of 1,381 (SE: 163, 95% CI: 1,097-1,738) individuals (Table 2). The model M(th), which assumed heterogeneous capture probabilities that varied with time, was the next most parsimonious model to explain the variation in the data according to the selection criteria value (0.76; Table in S3 Table). This model structure was also used by [21] to model abundance in Plettenberg Bay in 2002-2003. Because it is recommended that selection values lower than 0.75 should not be used to estimate abundance [35], comparison using this model was justified. The abundance estimate for this model M(th) was 1,494 (SE: 224, 95% CI: 1,131-2,024) marked individuals, giving a super-population estimate of 1,940 (SE: 291, 95% CI: 1,448-2,600) for the bay (Table 2). This is 72.3% lower than the estimate of 6,997 for Plettenberg Bay in 2002-2003 [21]. The closed population analyses for the 2014-2016 period were repeated using MARK (Table in S4 Table). The best model (based on ΔAIC) for Plettenberg Bay was p=c(t), denoting that the capture and recapture probabilities are equivalent and time dependent. This model predicted a super-population size for Plettenberg Bay of 1,386 (SE: 62; 95% CI: 922-2,083) individuals.

**Table 2:** Estimates of *T. aduncus* abundance based on closed population models conducted using CAPTURE in the entire study area, and for the Plettenberg Bay area in isolation for the periods 2002-2003 and 2014-2016. Estimate of marked population (N) and super-population size (NT); standard error (SE); lower and upper limits of the 95% confidence interval (LCL and UCL).

| Model <sup>1</sup> | Marked population |  |  |  | Super-population |  |  |  |
| --- | --- | --- | --- | --- | --- | --- | --- | --- |
|  | N | SE | LCL | UCL | NT | SE | LCL | UCL |
| Entire study area (2014-2016) |  |  |  |  |  |  |  |  |
| M(th) | 2103 | 144 | 1850 | 2417 | 2731 | 188 | 2387 | 3126 |
| Plettenberg Bay (2014-2016) |  |  |  |  |  |  |  |  |
| M(t) | 1063 | 125 | 858 | 1360 | 1381 | 163 | 1097 | 1738 |
| M(th) | 1494 | 224 | 1131 | 2024 | 1940 | 291 | 1448 | 2600 |
| Plettenberg Bay (2002-2003) <sup>2</sup> |  |  |  |  |  |  |  |  |
| M(th) | 4833 | 742 | 3612 | 6556 | 6997 | 742 | 5230 | 9492 |

<sup>1</sup> Model description: M(t) - time varying capture probability (p); M(h) - heterogeneous p; M(th) a combination of the above [35].
<sup>2</sup> Results extracted from [21].

## Discussion

The current lack of knowledge of *T. aduncus* abundance and trends in South Africa hampers conservation assessments [16]. This study contributes novel information to assist conservation management by reporting *T. aduncus* abundance estimates in the Agulhas Bioregion of the southern Cape, and apparent change in abundance for the Plettenberg Bay sub-region. The large number of identified individuals in this study (817) and the low re-encounter rates (27%) supports the notion that individuals observed within the study area are part of a much larger open population that ranges as far as Algoa Bay [20].

The best open population estimate for the entire study area gave super-population size estimate of 2,295 individuals. This estimate needs to be interpreted with some caution, because goodness-of-fit results suggested over-dispersion in the mark-recapture data, with strong transience and heterogeneity in capture probabilities between individuals. The most parsimonious closed population model {p=c(t)}, whereby the capture and recapture probability are equivalent and time dependent gave a similar estimate to that of the open model, namely 1,940 individuals (Table in S4 Table).

A closed population model was required to compare abundance estimates from the present study with estimates derived from data collected between 2002 and 2003 [21]. The comparison between the two study periods (more than 10 years apart) is important because there is no other information on changes in population abundance for this species in South African waters, leading to considerable uncertainty regarding the species’ conservation status [16]. The best estimate for Plettenberg Bay in 2002-2003, was 6,997 dolphins. In comparison, the two most reliable estimates for this study were 1,381 and 1,940 individuals. The latter estimate is 72.3% lower than the 2002-2003 estimate.

The low re-encounter rate of known individuals in the area may have been influenced by there being a sizable proportion of transient animals in the population. For future studies, this could potentially be remedied through greater search effort in the area. However this is often not realistic due to weather constrains and moreover it would imply exorbitant costs for the running of dedicated research vessels. Using the tourist vessels as platforms of opportunity is a possible alternative but there would have to be consistency in the methods used during searching and encounters. Another alternative for estimating abundance and monitoring change in the area is through aerial surveys using a distance sampling approach. Aerial surveys can cover much more ground in a day, but have disadvantages such as the need for almost perfect weather conditions and very good water clarity in order to have a good detection rate (e.g. when animals are underwater). Furthermore abundance estimates from aerial survey are likely to be negatively biased by only taking into account individuals that are in the study area at the time of the survey, whereas the mark-recapture open models allow for individuals to enter and leave the study area. Another important limitation of aerial surveys is undercounting bias whereby as much as two thirds of animals may not be detected during the surveys, as shown in previous aerial survey studies (e.g. [38]). For this reason it is recommended that if aerial surveys are used, twin platform surveys should be conducted (e.g. [39]) whereby two aircraft survey the same transect independently but minutes apart in order to estimate the number of missed sightings.

A pilot study consisting of nine aerial surveys was conducted during the study period, to test the practicality of surveying *T. aduncus* using this method [18]. Abundance estimates were not derived from aerial surveys because there were too few surveys (n= 9) for a robust population estimate. The group size estimates from boat surveys are, however, corroborated by the aerial survey estimates, with both survey methods detecting larger group sizes during winter [18].

The overall mean group size during aerial surveys along the entire study area was 43 ± 37 (range: 1-150; median: 33; n= 42), compared with 47 ± 55 individuals from boat-based surveys (Table 1). In winter, the estimate from aerial surveys was 46 ± 34 (range: 6-100; median: 39; n= 12) compared with 57 ± 63; and in summer, 41 ± 38 (range: 1-150; median: 30; n= 30) compared with 35 ± 42, respectively.

Smaller average group size were recorded in Plettenberg Bay (26 individuals) compared with the whole study area (47 individuals). Both these estimates are considerably lower than the mean group size of 120 that was estimated for 2002-2003 in Plettenberg Bay [21]; the decline in group size for Plettenberg Bay between the two periods was 78.3%. A decline in average group size is also corroborated by a shore-based estimate of mean group size from the early 1970s, of 140.3 [40]. The decline in group size may be an indication of a decline in numbers and appears to support the decline shown by the modelled abundance estimates.

Another factor that could have influenced the decline of group size and abundance is a reduction in the numbers of transient groups using the area. In several recent years South Africa’s annual sardine run which is characterized by large schools of sardine (*Sardinops sagax*) moving northwards along the east coast during winter months, followed by vast numbers of predators including *T. aduncus* [41], has been less pronounced than in the past [42]. The dwindling size of the sardine run could have the effect that less transient groups of *T. aduncus* navigate through the study area. Declines in the availability of other important prey resources for *T. aduncus* such as squid [43,44], which spawn in a distinct area around Plettenberg Bay [45] but which have been less productive in recent years[46] could also have affected *T. aduncus* numbers in the area.

An important change in Plettenberg Bay since the 2002-2003 study of *T. aduncus* is the growing resident Cape fur seal colony (*Arctocephalus pusillus pusillus*) on the Robberg Peninsula [47]. This could cause direct competition for prey resources with *T. aduncus* including for species such as: piggy (*Pomadasys olivaceum)*, squid *(Loligo vulgaris reynaudii)*, cuttlefish *(Sepia spp*.), red tjor-tjor (*Pagellus bellotii*), sardine (*Sardinops* sagax) and octopus (*Octopus spp*.) [43,48]. Furthermore, there is likely to have been an increase in the abundance of great white sharks (*Carcharodon carcharias*), that are attracted to seal colonies, in the area. This impact of the sharks on the *T. aduncus* population may be direct (i.e. predation in itself) [49]; or indirect, whereby the predation risk brings about increased stress levels in the prey population that can reduce their performance and productivity, or changes in residency patterns reducing time spent in the area [50].

Due to their coastal distribution, *T. aduncus* are also vulnerable to multifarious anthropogenic pressures associated with coastal and inshore areas that could bring about shifts in residency patterns or a population decline. In our study area such pressures include coastal development, vessel traffic and associated disturbance, especially those related with boat-based cetacean viewing ventures [6,51,52]. The longevity and relatively low reproductive rate of this species aggravates the effects of habitat degradation and other threats. The Bitou municipality (which includes Plettenberg Bay) is the fastest growing municipality in the Western Cape Province, with an average annual population growth of 4.8% from 2001 to 2013 and tourism brings in much revenue to the area [22]. However, while it may be tempting to link the decline in *T. aduncus* numbers and group sizes with the increasing population and associated pressures in the area, a considerable increase in the mean group size of the same species in the more developed Algoa Bay to the east has been shown, from 18 to 76 individuals between 2008 and 2016 [53], which can be a consequence of a shift of the population’s preferred habitat in recent years.

While the causes of the changes in Plettenberg Bay are not yet well understood, a precautionary approach especially with regard to impacts of the burgeoning tourism industry is advised, and this is naturally also in the interests of the industry’s sustainability. The impacts of tourism on animal populations is generally measured by short-term behavioural responses (e.g. [52]), yet evidence is mounting that disturbance caused by these activities have long-term demographic implications. In Plettenberg Bay, boat-based ecotourism may have impacted on the sympatric Indian Ocean humpback dolphins (*Sousa plumbea)*, which is known to be sensitive to human presence. Preliminary results have shown a decline in abundance of this population by approximately 46% between 2002-2003 [54] and 2012-2013 [55]. Simultaneously, a 35% reduction in the mean group size of this species between the two periods was documented [55].

In other parts of the world, *Tursiops spp.* have also been declining. For example, in Australia [56] and the Bahamas [57], declines of 15% and 49% were attributed to effects of tour operator vessels and a combination of natural and anthropogenic factors. Some of the measures that were taken in other parts of the world to mitigate impacts and protect *Tursiops spp.* includes the creation of protected areas (e.g. [58]).

While *T. aduncus* was recently assessed to be Near Threatened in South Africa [16], the *S. plumbea* is currently Endangered at the national level on account of the small size of the population and apparent decline, exacerbated by its fragmented distribution with considerable movement within the bioregions [59,60]. Expanding the current MPAs or identifying new conservation areas has been recommended for *S. plumbea* in South Africa [60]. Given the sympatry of the two species, such measures could also address certain conservation needs for *T. aduncus*; e.g. if vessel traffic is strictly controlled in such areas, if critical habitat types are protected and if human pressures on prey resources in such areas are reduced such that productivity and overspill of certain prey into adjoining areas may occur (e.g. [61,62]).

## Conclusions

This is the first study to show a change over time in abundance for the *T. aduncus* anywhere in South Africa. While a comparison based on closed population models between two periods for a population that is likely to be open in nature may not be ideal and intuitively should be accepted with caution, such a comparison was called for given the lack of such information on the species and resulting uncertainty regarding its conservation status in the country. Moreover, comparison of mean group sizes between the two periods 2002-2003 and 2014-2016 also showed a substantial decrease that corroborated the model-estimated decline in abundance during the same period. While the causes of the apparent changes are not yet well known, precautionary measures or controls to prevent and mitigate disturbance to the population and also that of the sympatric, Endangered *S. plumbea* are advised, especially with regard to potential disturbance associated with marine tourism activities. The results of this study highlight the need for further research and monitoring in the area as well as the importance of assessing abundance changes at other sites to inform revision of *T. aduncus* conservation status in South Africa.

## Acknowledgements

Special thanks to numerous individuals from Ocean Odyssey, Enrico’s Fishing Charters, Orca Foundation, Ocean Blue Adventures and South African National Parks (Tsitsikamma) for providing research vessels and skippers. Special thanks to students and volunteers who assisted with data collection. This research was conducted under a Memorandum of Understanding for research between Nelson Mandela University and the Department of Environmental Affairs (DEA), and would not have been possible without DEA support.

